# Neuroprotective potential of an *Ayurvedic* compound “Ras-Sindoor” on *Drosophila melanogaster* model of Parkinson’s disease

**DOI:** 10.1101/2022.11.18.517027

**Authors:** Sonia Narwal, Arushi Rai, Shreyas Iyer, Kirti Tare, Meghana Tare

## Abstract

Parkinson’s Disease (PD) is the second most common neurodegenerative disease affecting 1-2% of the global population with no cure to date. PD is characterized by accumulation of Lewy Bodies (LBs), which are caused due to aggregation of incorrectly folded α-Synuclein, (SNCA). Another form of PD manifestation is characterized by loss of function of *parkin*, which encodes an E3 ubiquitin ligase. Despite extensive research, the cause for onset and progression of PD remains unknown and current therapeutics mainly help manage the disease. An alternative line of treatment can be useful.

In this study, we have employed two different genetic models of *Drosophila* to screen for *Ayurvedic* compounds and found an *Ayurvedic* mercury based organo-metallic drug Ras-Sindoor has neuroprotective function. Our data indicate that characteristic locomotory dysfunction phenotype of PD is restored upon administration of the compound. Interestingly, RS fed flies also exhibit reduced transcript levels of initiator caspase *dronc*, which possibly prevents cell death in dopaminergic neurons. Additionally, RS fed PD model flies exhibit an enhanced life span. Our studies emphasize beneficial use of traditional *Ayurvedic* compounds as a holistic cure for PD like multifactorial diseases.

## Introduction

Parkinson’s Disease (PD) is the second most common age-related neurodegenerative disease after Alzheimer disease and affects the people with the prevalence of 0.8% of the worldwide population among the other neurological disorders[1][2]. The two major neuropathological features that characterize PD are: progressive loss of dopaminergic neurons in substantia nigra pars compacta region of mid brain, which affects the dopamine level in the striatum, and the presence of intra-neuronal Lewy bodies, which is mainly the aggregation of α-synuclein protein. With the disease progression, PD patients develop four major motor symptoms: tremor, bradykinesia, muscle rigidity, and postural instability [3][4][5]. Most of the PD cases are sporadic with unknown etiology, possibly environmental or genetic factors are involved in the development of the disease. About 5-10% cases of PD are familial and single gene mutations[6]. *α-synuclein* and *parkin* are two of the genes with mutations that have been linked to familial form of PD. Wild type and mutant form of α-synuclein (A53T and A30P), which is a presynaptic membrane protein reportedly involved in formation and release of synaptic vesicle, have been found to cause the early-onset autosomal dominant PD[7][8]. Mutations in *parkin* gene, which encodes an E3 ubiquitin ligase, have been found to be associated with autosomal recessive PD[9]. Apoptosis, mitochondrial dysfunction, oxidative stress due to α-Synuclein accumulation and *parkin* mutation have been reported to be major causes for loss of DA neurons [10][11][12] .

To date, there is no cure or effective therapies to slow or stop the progression of PD. Although there are several drugs such as levodopa, bromocriptine, pramipexole and ropinirole; MAO-B inhibitors, such as rasagiline; catechol-O-methyl transferase (COMT) inhibitors such as entacapone and selegiline are used to alleviate the motor symptoms only and cause several side effects[13][14][15]. Therefore, identification of new therapeutic strategies is required that can target PD symptoms and slow down the disease progression.

*Ayurveda* “The Science of Life” is the world’s oldest traditional system of medicine that originated in India around 5000BCE [16]. *Ayurveda* medicinal system also claims to provide healthy aging and is reported to alleviate neurodegenerative disorder symptoms[17][18][19]. The classical manuscripts describing Rasayanas and Bhasmas form a branch of pharmacology named Rasa-shastra. Rasa-shastra broadly can be categorised into two medicinal forms viz. Bhasmas of Minerals and their combinations with herbs, and Ras-kalpa known as Rasayanas. Further, they are classified as Paarad/mercury based and non- mercury-based preparations, depending on the inclusion of Paarad/mercury and its sulphide, Gandhak. As per the method of preparation, they are categorised as Kupipakwa Rasayan (Paarad-Gandhak combination when heated at a certain temperature in a Puta/furnace). This is performed to obtain the medicine in the form of sulphides of mercury. The next type is Kajjali (Paarad-Gandhak), which is processed together in a mortar and pestle, named as Kharaliya Rasayan. Few examples of Rasayanas are Ras-Sindur, Siddha Makardhwaja, Bruhadvaat Chintamani Ras.

Bhasmas like Rajat Bhasma, Suvarna Bhasam, Sahasraputi Abhrak Bhasma andTamra Bhasma are prepared by combining with different herbs for their purification, till the formation of Bhasma after heating it at a defined temperature in a furnace. Process of the formation of Bhasma is known as bhasmikarana. In *bhasmikarana* process, a metal is converted into a desired chemical compound through incineration with the use of some minerals or herbal extract to neutralize the toxicity of metal. Further, *Bhasma* is subjected to physico-chemical processing called *Samskaras* to purify, detoxify and retain the therapeutic properties[20]. One of the mercury based compounds, Ras-Sindoor (RS) is a well-known *Ayurvedic* drug in the form of mercury sulphide (Hg-S), with crystalline size of 25-50nm [21]. It has been shown that RS promotes healthy aging and suppresses the neurodegeneration in *Drosophila melanogaster* model of Alzheimer’s Disease (AD) and Huntington’s Disease (HD) [22].

Apoptosis, a process for programmed cell death, is a part of developmental process and a response to any physical, genetic, and metabolic stress like oxidative stress, trophic factor deprivation, excitotoxicity of calcium ions. Neurodegeneration majorly involves death of neurons by apoptosis and formation of apoptotic bodies. These apoptotic bodies are further removed by phagocytosis. Protein degradation and caspases activation are the major hallmarks of apoptosis. The biochemical components of apoptotic pathways include groups of B-cell lymphoma (Bcl-2) family, apoptotic peptidase activating factor (Apaf-1), and caspases [23].

Caspases are inactive zymogen molecules which get cleaved to activated form in response to stimuli like death signal, stress. There are 3 major classes: inflammatory caspases involved in inflammatory responses to microbial interactions like caspase 1, 4; the apoptotic initiator caspases regulate the activation of downstream caspases to progress with a caspase activation domain as present in caspase 9, or with a death effector domain like in caspase 8, 10; the third type of caspase is the executioner caspases with a short pro-domain and get activated after cleavage process regulated by initiator caspases. The death effector domain caspases are a part of extrinsic apoptosis and intrinsic apoptosis is mediated by caspase activation domain consisting of initiator caspases-caspase 9. Previous reports suggest strong relation between intrinsic apoptosis, mitochondria related apoptosis and Parkinson disease progression [23].

In this study, we have used *Drosophila melanogaster* model of PD to test the efficacy of RS in alleviating the PD symptoms. *Drosophila melanogaster* model organism has several advantages including a higher degree of conservation in developmental pathways with mammals including apoptosis [24]. Transgenic *Drosophila* has been extensively used as a model system to study several neurodegenerative diseases: Amyotrophic Lateral Sclerosis (ALS), Huntington’s Disease, PolyQ Disease [25][26][27] and many other developmental disorders [28]. This model has also been used to study therapeutic effects of various natural compounds [29]. *Drosophila* as model system for PD, also recapitulates the major neuropathological features of PD, e.g. Lewy Body formation, locomotory defect and loss of Dopaminergic neurons [30] [31].

Here, we show that RS alleviates the locomotory defects as well as decreased survival rates in the flies exhibiting neurodegeneration induced by either over-expressing wild type or mutated *α-synuclein*; and, upon downregulation of the *parkin* in a dose dependent manner. We also identify that RS suppresses the initiator caspase *dronc* at a transcriptional level in the PD exhibiting flies from an early developmental stage. Our studies thus confirm a neuroprotective function of RS from an early developmental stage.

## Material and methods

### Fly Stocks

*Actin Gal4; Ubi GFP/TM6bTB* (a gift from Lakhotia lab, BHU, India), *w^118;^ Elav-Gal4, w+; +; Ple Gal4* (BL-8848),*UAS-Mito GFP/CyO* (BL-8842), *UAS-SNCA (A30P)* (BL-8147), *UAS-SNCA* (51375),*UAS-Park RNAi/ CyO* (BL-37509), *UAS-Park RNAi* (BL-31259) fly lines were used. All the stocks were maintained on standard fly food containing agar, maize powder, yeast and sugar at 25°C. Using Gal4/UAS system appropriate genetic crosses were carried out to obtain desired genotypes.

### Ras-Sindoor (RS) feeding

Genetic crosses were set up on egg collection medium containing agar, sugar, and fruit juice. Eggs were collected and transferred to the standard fly food and food containing different concentrations (weight/volume) of Ras-Sindoor i.e., 1%, 0.5%, 0.25% and progenies from these crosses were used for the desired experiments. For each experiment, the regular and the RS containing food were prepared from the same batch; similarly, all adults for an experiment were derived from a common pool of eggs of the desired genotype and fed in parallel on the regular or the RS food. RS was commercially prepared by Arya Vaidya Sala, Kottakkal, Kerala, India. All the crosses were maintained at 29 °C.

### SEM-EDS sample preparation and analysis

RS fed flies were collected into cryo-vials and snap frozen using liquid nitrogen. Frozen flies were transferred into microcentrifuge tube containing 500ul of ice-cold acetonitrile (50%) and homogenized using homogenizer. Homogenized samples were then centrifuged at 12,000 rpm for 10min at 4°C. Supernatant was collected into pre-weighed microcentrifuge tube and lyophilized using speed-vac vacuum concentrator.

Lyophilized sample was then subject to SEM-EDS analysis. For EDS-analysis, samples were mounted on a metallic stub using a carbon tape and sputter coated with gold metal using Q150T ES Quorum system. SEM analysis was carried out in a Thermo Scientific Apreo S SEM system. EDS analysis was performed using Oxford instrument X-Max-N at 20 kV.

### Survival Assays

For the survival assays, freshly eclosed flies of desired genotypes, from regular food and RS containing food, were collected into respective food vials. Flies were transferred to new fresh food vials every other day without anesthetization and the number of dead & surviving flies were recorded each day until the desired age (30 days). Three replicates were carried for each genotype and a percentage (%) of survival were calculated. Flies were maintained at 25°C.

### Locomotor Activity

To characterize the locomotory dysfunction, climbing assays were performed. Flies were aged up to 30 days in the regular and RS containing food. Groups of 10 flies per genotype were transferred into cylindrical glass tube after anesthetization and left for 5-10 min for the revival and acclimatisation at room temperature. Tubes were marked at 8cm above the bottom of the vial. After acclimatization, the flies were gently tapped down to the bottom of vial and the number of flies that crossed the 8cm mark were recorded after 10 sec. Three trials were performed, and numbers were then averaged, and the resulting mean was used as the overall value for each single group of flies. For all genotypes, three replicates were carried out.

### Quantitative Real-Time PCR

Total RNA was prepared from adult fly or heads using the TRIzol method (Invitrogen) referred from Jove protocol (URL: https://www.jove.com/video/50245). cDNA was synthesized using Verso cDNA Synthesis Kit (AB1453A). RP49 was used as reference gene to normalize the Dronc amplicon. Quantitative RT-PCR was performed using 5x HOT FIREPol^®^ EvaGreen^®^ qPCR Mix Plus (Solis BioDyne). Primer Sequences were as follows:

*RP49*: Forward:5’-CCA AGG ACT TCA TCC GCC ACC-3’

Reverse: 5’- GCG GGT GCG CTT GTT CGA TCC-3’

*Dronc*: Forward: 5’-CTCGCTAAACGAACGGAGAAC-3’

Reverse: 5’-CAACGACACCCACATAAGGG-3’

### Statistical Analysis

Statistical analysis were performed using GraphPad Prism 8.0.1 software. Data are expressed as mean with SEM of replicates. The p-value and statistical significance are as per legends.

## Results

### Ras-Sindoor (RS) feeding has no toxic effect on transgenic *Drosophila melanogaster* model of PD

To test the non-toxic effect of RS in flies, the transgenic flies were fed with selected concentrations of RS and maintained for the time points required for assays and experiments. RS contains mercury in the mercury sulphide form (HgS) and its elemental and structural characterization has been well documented which shows that it is chemically pure α-HgS with Hg: S ratio as 1:1 [21]. The toxicity was tested based on the observation if the stable 1:1 Hg:S is altered after being consumed by the flies. So, we performed the EDS analysis of lyophilized Ras-Sindoor from RS fed flies and pure RS powder (RS from Kottakkal Ayurveda) as control. In EDS analysis, we found that element % and atomic % of Hg and S from fed flies (Fig 1c-d) were same as with RS powder form (Fig 1a-b). In both the samples, atom balance shows that both Hg and S are present. Hence, we confirm that due to presence of native configuration of Hg:S form in the RS fed flies with respect to pure RS powder, the compound causes no toxicity to the flies after feeding.

**Figure 1:**
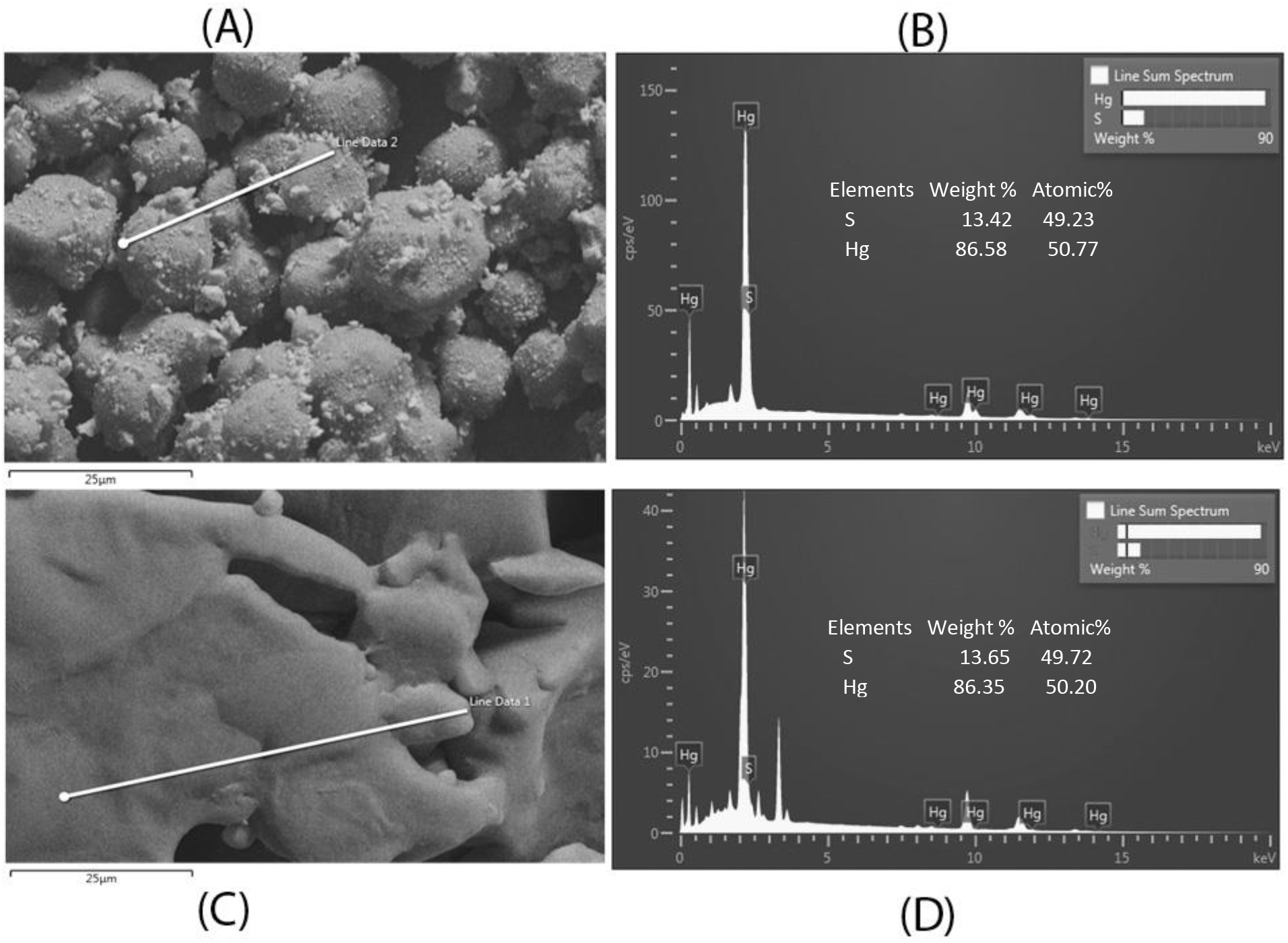
SEM-EDS analysis of Ras-Sindoor.SEM micrograph and EDS Spectrum of Ras-Sindoor (A, B). SEM micrograph and EDS Spectrum of Ras-Sindoor from fed flies (C,D).

### Ras-Sindoor (RS) rescues locomotor defects in PD flies

After confirming the non-toxicity of RS, we assessed the effect of RS on PD flies. First we established the locomotor dysfunction in PD flies using different GAL-4 lines to either overexpress *α-synuclein*, or to downregulate *parkin* (using UAS *parkin* ^RNAi^), at different time points (7 days and one month), as PD is an age and dosage dependent disorder. The flies were cultured from the embryonic stage to adult stage on RS supplemented food with three different concentrations: 1%, 0.5%, 0.25% (w/v). We performed locomotory assays of one week and one-month old RS Fed flies. We found that, in the ubiquitous *parkin* downregulation and *SNCA* overexpression model (Actin GAL-4), enhanced climbing ability for the concentration 1% RS in the 7-day time point, i.e. flies aged till 7-days fed on RS was observed. The 0.5% and 0.25% too had enhanced ability but to a lesser extent as to what was visible in 1% RS fed flies. When these flies were aged till 30 days, the climbing assay was performed again, and 0.25% and 0.5% RS fed flies were found to have increased climbing ability as compared 1% RS fed flies (Fig 2a-b). When we downregulated *parkin*, or overexpressed *SNCA*, pan-neuronally, and fed with the same concentrations of RS, the 1% RS fed flies exhibited enhanced locomotory behaviour as of 7-day old state. But this pattern turned in favour of 0.5% and 0.25% RS eventually at 30-day old stage, owing to the plausible reason of higher concentration RS toxicity (Fig 2c-d). When the above groups and feeding protocol were repeated and expressed specifically in dopaminergic neurons tissues, flies depicted significantly enhanced climbing ability when fed with 1% RS as compared to other groups in all RS fed and control (Fig 2e-f).

**Figure 2.**
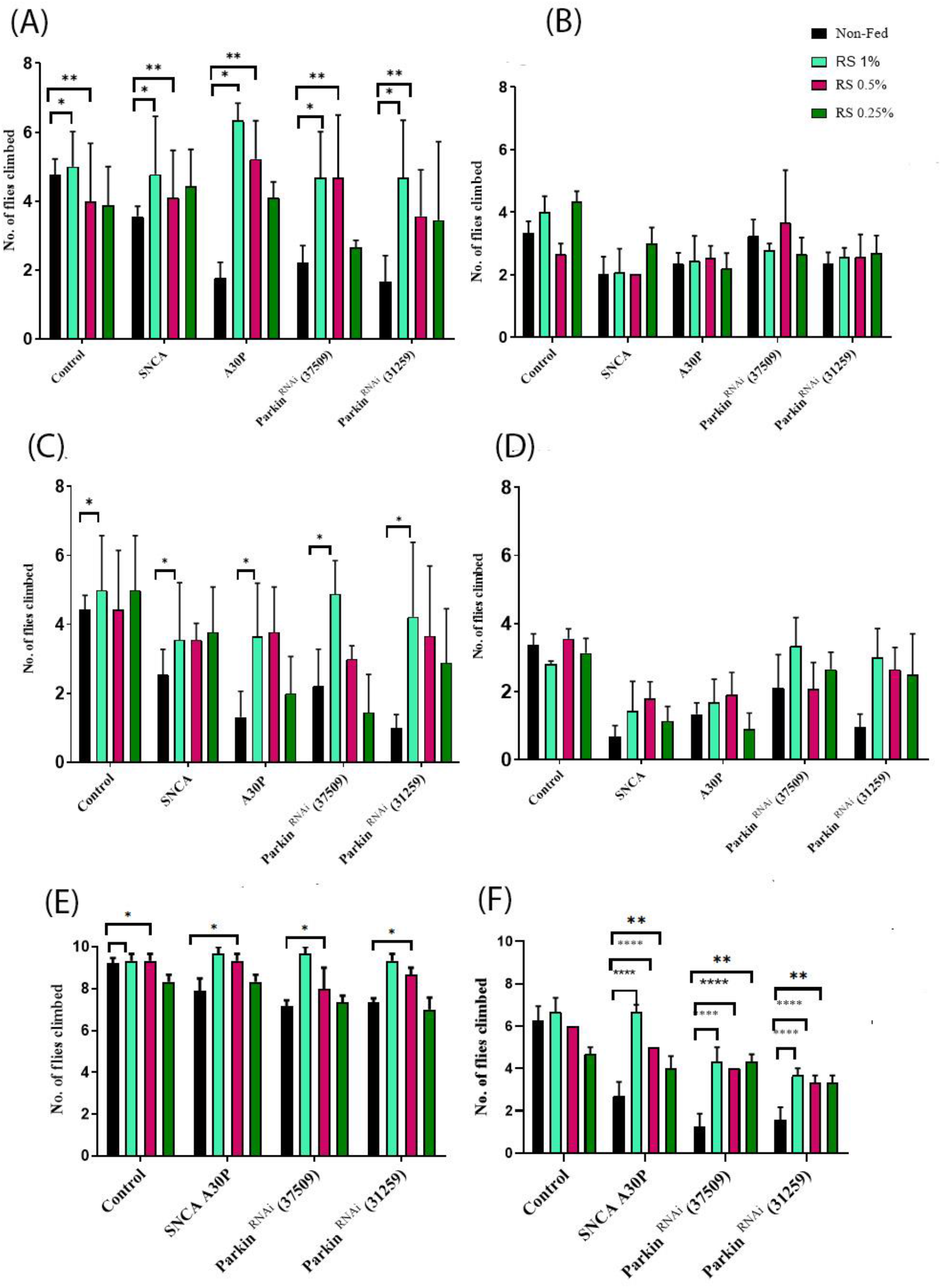
Improved locomotion of one week and one-month old of RS fed versus non-fed flies. (A,B) Actin-Gal4. (C-D) Elav-Gal4. (E-F) Th-Gal4. Graphs represented in A, C, E are of one week old flies, whereas B, D and F are from one month old flies. Data is represented as mean with SEM. (Two-way ANOVA followed by Dunnett’s multiple comparison test. P value: *(.033), ** (.002), *** (<0.001).

### Ras-Sindoor (RS) feeding improves life span of the PD flies

PD patients have reduced life span after diagnosis. Mainly, life span has been found to be more reduced in early onset (at the age of 50 years) as compared to late onset (at the age of 70 years) PD patients [32][33]. With the above experiments involving climbing assays, we observed consistent improvement in the performance of flies after being fed with RS. So, we wanted to understand the effect RS has on viability of the flies and whether this effect is in synchronisation with the improved climbing ability. Hence, we also tested the survivability of the RS fed and non-fed flies for one month. Survival rate of RS fed flies, expressing *α-synuclein* and *parkin RNAi* ubiquitously (Fig 3a) and in neuronal cells (Fig 3b), were found to be elevated as compared to non-fed flies. The rescue in viability was higher in case of RS feeding in ubiquitous condition as compared to neuron specific condition.

**Figure 3.**
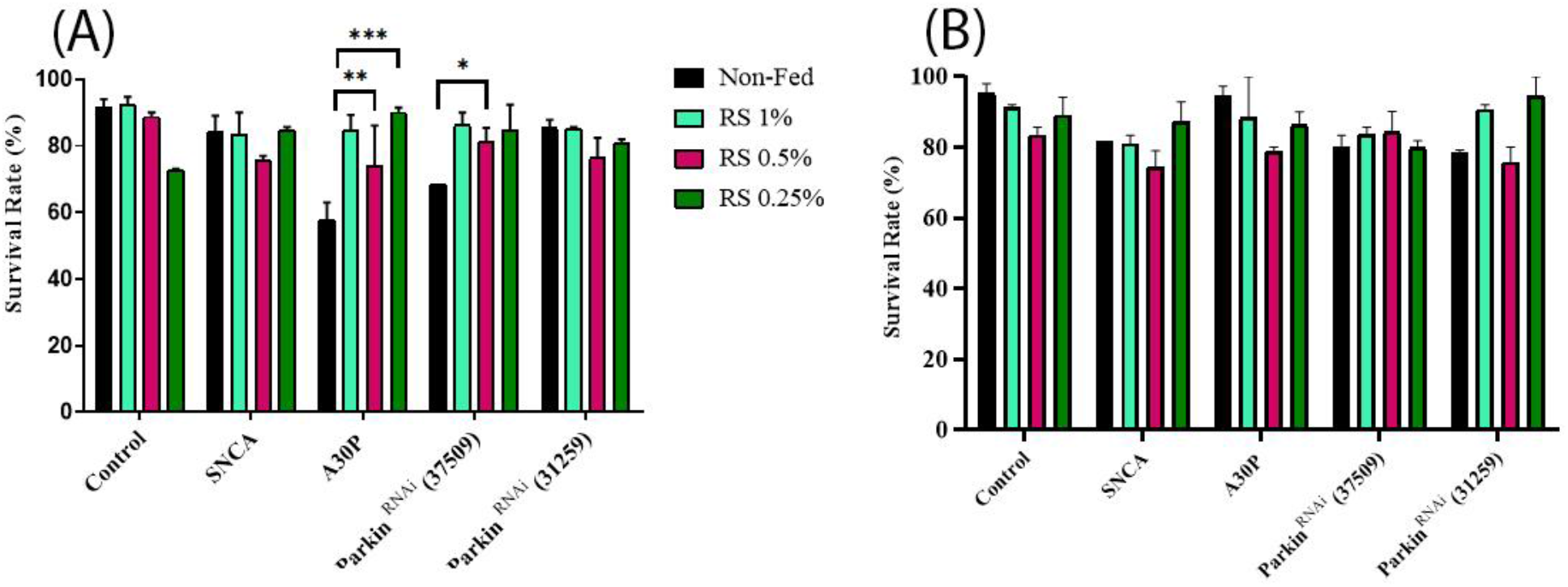
RS feeding in Parkinson’s Disease flies improves survival over a period of time. (A) Actin-GAl4. (B) Elav-Gal4. Data are represented as mean with SEM. (Two-way ANOVA followed by Dunnett’s multiple comparison test. P value: *(.033), **(.002), ***(<0.001).

### Ras-Sindoor feeding causes reduced apoptosis in Drosophila model of PD

One of the known mechanisms for loss of DA neurons is apoptosis. Intrinsic apoptosis has been considered more crucial for PD progression; thus, we have tested *Dronc* expression levels. We hypothesized that improved climbing ability could be due to decreased apoptosis in PD exhibiting flies upon RS feeding. We observed enhanced climbing ability and viability in flies fed with RS for a duration of one month. We therefore tested the transcription level of initiator caspase *dronc* in RS fed flies versus non-fed flies through real-time PCR using total RNA isolated from the whole flies expressing either *α-synuclein*, or *SNCA A30P* or the flies with downregulation of *parkin* in ubiquitous manner using *Actin-Gal4* transgene. Interestingly, we found that *dronc* transcriptional level was significantly reduced in RS fed flies as compared to non-fed flies (Fig 4).

**Figure 4.**
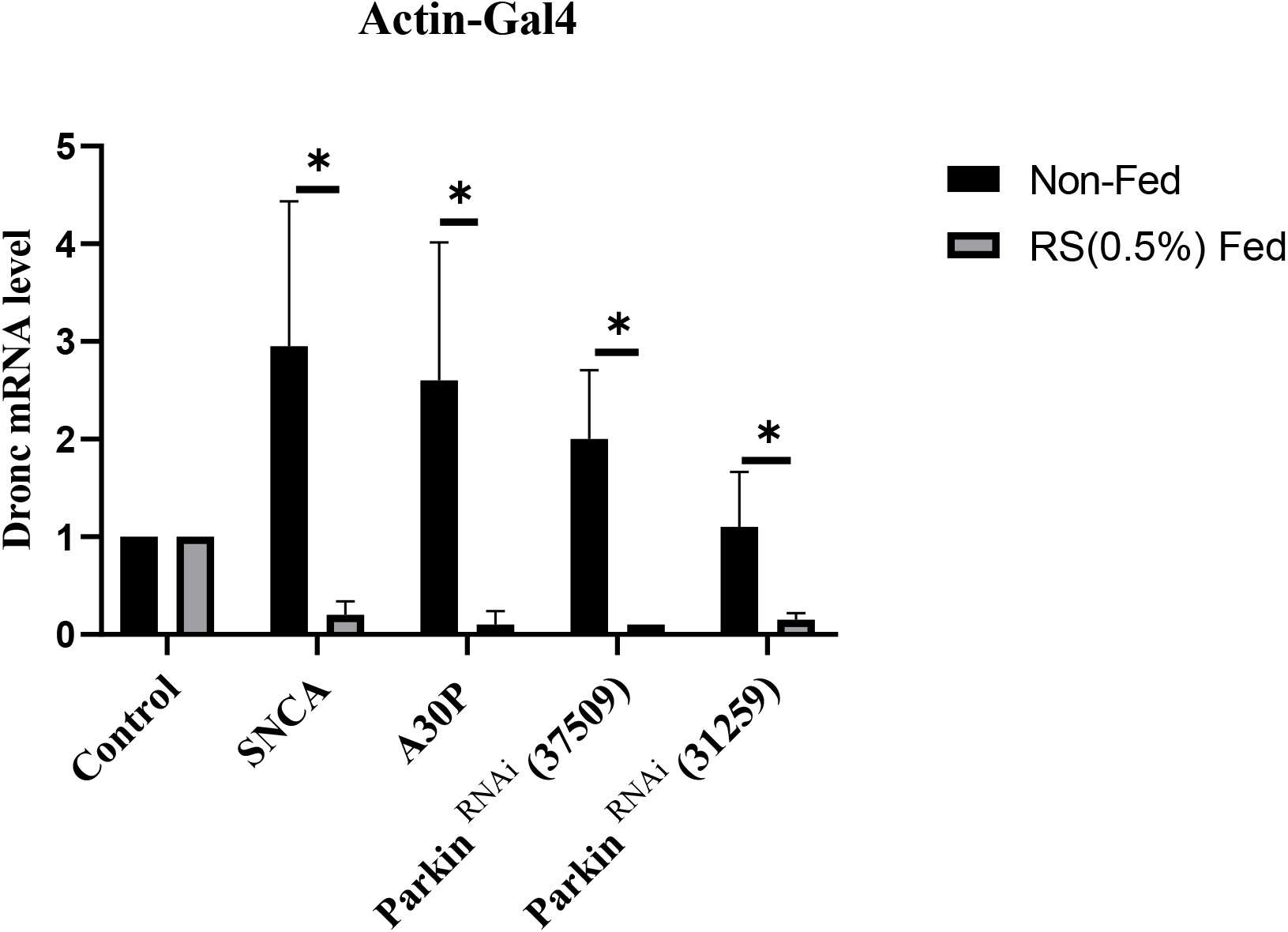
RS feeding results in reduced transcription of caspase *dronc*. two-tailed t-test (*P<0.05).

## Discussion

Parkinson’s Disease, or the shaking palsy, results into loss of dopaminergic neurons and causes locomotory defects in the patients. The evolving trends in the field of therapeutics are taken from nature, which has always been a great support to all the life, including humans who exhaust the nature of all its resources. There have been solutions implemented for this age dependent neurodegenerative condition, but these methods pose certain side effects or drawbacks. COMT inhibitors reduce dopamine metabolism in the body but do not affect non-motor symptoms [13]. Levadopa causes dyskinesia after intital improvement in the motor symptoms [34]. Deep brain simulation is beneficial in motor symptom treatment but poses adverse effects sometimes in speech impairment. Various other cell based therapies are also implemented but they are not reliable due to low graft survival, ethical issues. Viral vector mediated gene delivery has also been useful but with limitations of size limit of delivery, risk of mutation. Hence, considering the pros and cons of selected present therapeutic techniques, utilization of natural compounds can be a very reliable [13]. *Ayurveda* is a traditional medicinal system being practised sinceages. It mainly works by the metabolites and products derived from plants and mixed with or stabilized with certain specific inorganic components [34]. There are studies highlighting the neuroprotective effects of various other natural compounds in terms of PD in various models of study like in mice exhibiting inhibition of MAO-A activity in the brain by *Curcumin longa*, or increase of neurotropic factor expression like BDNF in rats on exposure to curcumin. When human patients were fed with cooked *Vica faba*, they expressed improved clinical PD signs, enhanced plasma levels of L-dopa, increased motor response duration [35].

There are considerable amount of literature supporting the therapeutic potential of Ras-Sindoor (RS). Hence, we wanted to make sure there was no toxicity induced due to mercury or sulphide, in the Mercury sulphide complex. Based on the method of preparation of RS, it is quite clear that the compound is thermostable even at extreme temperatures [21]. Hence, when analysed for the difference in the atomic % and elemental % of Hg and S of the RS compound in its native form and after ingestion by the flies. We performed the EDS analysis and found the same percentage in ingested form as compared to native state. These results are not surprising, since there is an existing report discussing stability of RS using XRD, XPS and TEM. The thermal stability is also high up to 344°C melting point. Due to its non-toxic, nano-crystalline nature with increased surface area, this study has also pointed out the potential of RS as a vector in delivering drugs [21].

Since, we have established that the RS has no toxic effects on the health of the living organisms, we further observed the effects of RS feeding on the motor symptoms in PD conditions. Hence, the flies when fed with RS different concentrations, 1%, 0.5% and 0.25%, with transgenic mutation expressed ubiquitously, in neurons and specifically in dopaminergic neurons, the climbing ability of these flies were assessed and found to be rescued. The significance was higher only in 0.5% RS fed flies in 7-day old ubiquitous *SNCA* overexpression and *parkin* downregulation flies. These data are supported by another similar report, where, concentration of RS at 0.5%, was found to be more effective on the viability of the model organism [22]. The concentration of 0.5% RS is also preferred for the same reason [34]. This rescue was not significant in many conditions such as, one month old ubiquitously expressed, or/and pan neuronal *SNCA* overexpression and *parkin* downregulation flies, or 0.25% RS fed conditions on 7-day old flies. This may be because the flies would be requiring to age more to exhibit more deterioration and then rescue of the climbing ability due to PD progression. However, when flies expressing *SNCA* overexpression and *parkin* downregulation specifically in dopaminergic neurons, were fed with the 3 concentrations of RS, all three showed significant rescue of climbing ability in one month old flies. This might be due to the higher effect of the *SNCA* overexpression and *parkin* downregulation on the organism due to direct activation in the dopaminergic neurons only. Hence, when fed with RS, the rescue is also significant. This model confirms the efficacy of RS as a therapeutic compound for PD. The data obtained here for RS compound is one of the many natural compounds found to have neuroprotective effects. Another study has reported rescue of locomotory defects caused due to *PINK-1* mutation in flies, upon feeding with extracts of *Zandopa, Withania somnifera, Centella asiatica* [36].

PD is an age dependent neurodegenerative condition and is found to severely affect the life span if detected in early life (before 70 years). Hence, the number of years lived with PD is low [37]. Survival assays were performed on mutant flies fed with respective RS concentrations, and it was evident that when *SNCA* overexpression and *parkin* downregulation expressed ubiquitously, the survival rate of flies fed with 1% RS was higher as of other groups in each treatment, including control. But this difference was less and non-significant when flies expressing these pan-neuronally were analyzed for the same. The significance of rescue of viability was more in case of mutated *SNCA* overexpressing flies ubiquitously. This might be due to the severity of mutation since expressed ubiquitously, hence affecting the life span and degree of rescue as in case of *SNCA A30P* was higher. These data are in agreement with works that highlight the increased survival, fecundity, stress tolerance when fed with RS at above mentioned concentrations[34]. There are many other studies that discuss and explore the therapeutic and neuroprotective role of various classes of natural compounds extracted from plants like *Curcumin longa* has been reported to enhance viability of cortical neuron cells and downregulating *NFκB* transcription and ROS intracellular accumulation [38].

Neurodegeneration has been proposed to occur due to excessive apoptosis activation in the affected cells as observed in models of PD, AD. Intrinsic apoptosis has been found to be predominant in PD. Various lines of evidence suggest initiator *caspase 9* is known to play important role in intrinsic apoptosis and therefore, we tested the transcription level of *Dronc* (fly homologue for *caspase 9*), after feeding the *SNCA* overexpressed and *parkin* downregulated flies with 0.5% RS in the food [39]. This stemmed from the consistent improvement shown by flies in survival and locomotory assays after being fed with 0.5% RS and reports published before [34]. Ubiquitous expression system was selected for understanding the effects of the *SNCA* overexpression and *parkin* downregulation in the whole fly body and testing for the level of *dronc* transcript. Significantly reduced *dronc* transcription level were found in *SNCA* overexpressed and *parkin* downregulated flies after being fed with RS 0.5%. However, this confirms the role of RS as a neuroprotective agent against PD in transgenic models involving *SNCA* and *Parkin* and modulate cell death pathway. There are many other natural compounds which have been used as a therapy for PD and found to have anti-apoptotic effects in cell lines or model systems. This includes *Ginseng, Ginkgo biloba, Hypericum perforatum*, in repsonse to various neurotoxin induced PD by inibitng *caspase 3* activity, *bax*, *bcl2* increase [40].

## Conclusion

Parkinson’s disease is known to be mainly regulated by altered functions of *α-synuclein* and *parkin*, but other players like *dLRRK, DJ-1* have also been involved. It is a neurodegenerative condition that progresses slowly with age. There are several studies reporting many therapeutic strategies using chemicals and natural compounds from plants. It has been observed that the side effects using natural compounds are relatively less, and hence they are proving to be reliable. *Ayurveda* being a traditionally followed medicinal system is observed to rescue defects in PD conditions, as evident from ours and previously reported studies, but the exact mechanism of action and interacting partners is not known. RS has been shown to be effective in treating infections, and immune disorders. Our data indicates enhanced locomotion and survival of PD exhibiting transgenic fly models involving *SNCA* and *Parkin*. We relate these restorations to decreased levels of caspase-9, hence highlighting possible involvement of apoptotic regulation in RS therapy. Based on these observations and previously published data, RS is a reliable potential therapeutic agent against PD (Fig. 5). Further validation in other model organisms for molecular targets is required for substantiating the role of RS in Parkinson’s spectrum neurodegenerative disorders.

**Fig. 5:**
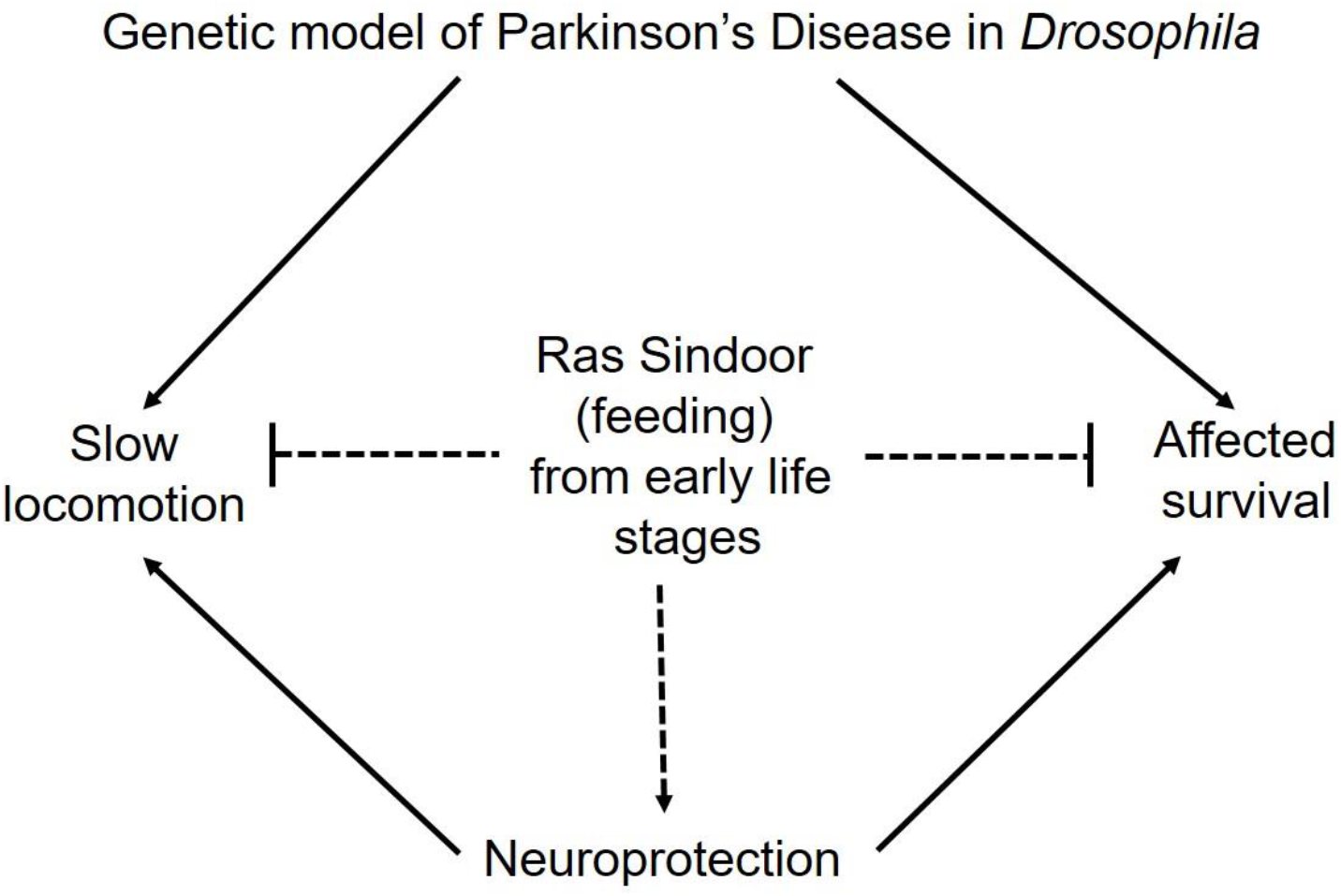
Potential therapeutic effects of Ras Sindoor on phenotypes exhibited in Parkinson’s Disease.

## Acknowledgement

Authors acknowledge Prof. Lakhotia for inputs on experiments, fly community, India for providing reagents, FlyDakia for procuring fly stocks, Kotakkal arya Vaidya shala for Ras Sindoor. Part of this work is supported by research initiation grant and additional competitive research grant to MT. SMI is funded by DST/SERB. SN is funded by BITS Pilani, Pilani.

## Author Contributions

MT conceptualized the work, KT provided insights for understanding approaches to use Ayurvedic compounds. SN, SMI and AR performed the experiments. SN and MT prepared the manuscript.

